# Targeted enzyme assisted chemotherapy (TEAC) – a novel microRNA-guided and selenium-based regimen to specifically eradicate hepatocellular carcinoma

**DOI:** 10.1101/2021.02.09.430426

**Authors:** Arun Kumar Selvam, Rim Jawad, Roberto Gramignoli, Adnane Achour, Hugh Salter, Mikael Björnstedt

**Author notes:** Correspondence to: Mikael Björnstedt.

## Abstract

Despite progress in the treatment of non-visceral malignancies, the prognosis remains poor for malignancies of visceral organs and novel therapeutic approaches are urgently required. Here we introduce a novel therapeutic regimen by treatment with Se-methylselenocysteine (MSC) and concomitant tumor-specific induction of Kynurenine aminotransferase 1 (KYAT1) in hepatocellular carcinoma (HCC) cell lines, using either vector-based and/or lipid nanoparticle-mediated delivery of mRNA. Supplementation of MSC in KYAT1 overexpressed cells resulted in significantly increased cytotoxicity as compared to MSC alone. Furthermore, microRNA antisense targeted sites for miR122, known to be widely expressed in normal hepatocytes whilst downregulated in hepatocellular carcinoma, were added to specifically limit cytotoxicity in HCC cells, thereby limiting off-target effects. KYAT1 expression was significantly reduced in cells with high levels of miR122 supporting the concept of miR-guided induction of tumor-specific cytotoxicity. The addition of alpha-ketoacid favored the production of methylselenol, enhancing the cytotoxic efficacy of MSC in HCC cells, with no effects on primary human hepatocytes. Altogether, the proposed regimen offers great potential to safely and specifically target hepatic tumors that are currently untreatable.

## Introduction

Combinations of first- and second-generation cytostatic drugs, novel targeted therapies including checkpoint inhibitors, multi-kinase inhibitors and immunotherapies have resulted in remarkable progress in the treatment of non-visceral malignancies, including breast cancer, renal cancer and malignant melanoma (1–3). However, the prognosis remains poor for visceral malignancies *e.g*. liver, bile duct and pancreas cancer, where most efforts so far have led to no or only marginal efficacy. Selenium (Se) compounds have been recognized for their cytotoxic properties, especially in tumor cells resistant to standard cytostatic regimens. The therapeutic potential in humans has not yet been demonstrated due to very few systematic human clinical trials and the lack of tools to increase tumor specificity.

Methylseleno-cysteine (MSC) is a natural selenium compound produced by plants as a detoxifying pathway (4). MSC may be considered a prodrug since it is inert and non-toxic in the absence of metabolizing enzymes. MSC has high bioavailability in humans and displays favorable pharmacokinetic properties with a short half-life and low risk for chronic selenosis (5). In mammals, MSC is metabolized primarily by kynurenine aminotransferase1 (KYAT1), a multifunctional, pyridoxal 5′-phosphate (PLP)-dependent enzyme. KYAT1 belongs to a class of enzymes that cleave carbon-sulphur (C-S) bonds with multiple substrate specificities (6). This cytosolic enzyme displays dual activities, transamination and β-elimination on single substrates. The nature of the substrate and the concentration of α-ketoacids determine which pathway will prevail, transamination or β-elimination. KYAT1 has also been shown to effectively cleave C-Se bonds in addition to C-S bonds (7), which renders MSC a suitable substrate for KYAT1. KYAT1 metabolizes the relatively non-toxic MSC through either transamination or β-elimination to β-methylselenopyruvate (MSP) (8) or methylselenol (MS) (9), respectively. Both MS and MSP have profound anti-tumor and anti-proliferative properties (10). While the tumoricidal activity of MSC has been previously demonstrated in several human cancer cell lines and animal models (11–13), MS is one of the most reactive metabolites of selenium and causes oxidative stress with pronounced ROS production and depletion of thiols and NADPH in cells (14, 15). A limited amount of studies indicate that MSP acts as a histone deacetylase (HDAC) inhibitor, leading to increased expression of tumor-suppressor genes (8). Hence, the selective induction of KYAT1 in tumor cells in the presence of MSC may represent an attractive route to selectively induce cytotoxicity in tumor cells.

RNA-based therapeutics allow the efficient transient induction of proteins via the delivery of exogenous mRNA into targeted cells or tissues (16). The benefit of RNA-based delivery over conventional DNA-based delivery is that mRNA does not integrate into the host genome. Protein translation occurs in the cytoplasm; hence mRNA-based delivery results in rapid and efficient protein expression in tissues (17). The main challenge in RNA-based therapeutics is to selectively induce the protein of interest into the targeted tissues and so reduce off-target cell protein expression. MicroRNAs are non-coding small RNA molecules frequently described as tissue- and disease-specific (18). Jain *et al* demonstrated that the inclusion of microRNA antisense target sites (miRts) into the 3’ UTR of mRNAs reduces the expression of introduced mRNA via endogenous microRNA in normal tissues. MicroRNA122 is known to be widely expressed in normal hepatocytes whilst downregulated in HCC. The liver-specific microRNA miR122 accounts for about 70% of total microRNAs in this organ and is important for liver metabolism and hepatocyte differentiation (19). The same study demonstrated that miR122 deletion resulted in epithelial-mesenchymal transition (EMT) and spontaneous HCC development in a miR122^−/−^ mouse model (19).

In the present study, we aimed to potentiate MSC cytotoxicity in HCC cell lines by inducing KYAT1 expression via mRNA transfection. We also aimed at mitigating off-target cell protein expression by incorporating microRNA antisense targeted sites (miRts) specific for normal hepatocytes. Altogether, the presented results indicate that exogenous delivery of LNP-encapsulated KYAT1-encoding mRNA, alongside the pharmacological elevation of MSC levels, may represent a compelling novel strategy to efficiently treat HCC.

## Materials and Methods

### Chemicals and reagents

Potassium dihydrogen phosphate, Di-potassium hydrogen phosphate, EDTA, Sodium hydroxide, Aminooxyacetic acid (AOAA), L-Tryptophan (L-Try), 3-Indoleacetic Acid (IAA), Indole-3-pyruvic acid (IPA), Phenylpyruvic acid (PPA), Homoserine (HS), DL-Propargylglycine (PAG), α-Keto-γ-(methylthio) butyric acid sodium salt (KMB), se-Methylselenocysteine hydrochloride (MSC), Dimethyl-2-oxoglutarate (α-KG), L-Phenyl alanine (L-Phe), 2-Amino-2-methyl-1,3-propanediol, Pyridoxal 5’-phosphate hydrate (PLP), 2,4-Dinitrophenylhydrazine (DNPH), Phenylmethanesulfonyl fluoride (PMSF), RIPA buffer, Trypan Blue, Protease inhibitor cocktail mix, N-N-Dimethyl formamide and pLKO.1 vector was purchased from Sigma-Aldrich (St.Louis, MO, USA). BFF-122 was purchased from Axon Medchem (Groningen, Netherlands), NADPH was purchased from Acros Organics (Geel, Belgium). Lipofectamine 3000 was purchased from Invitrogen (Invitrogen, Camarillo, CA, USA). Page Ruler Plus Prestained protein ladder was purchased from Thermo Fischer Scientific (Rockford, IL, USA). Phusion DNA polymerase, FastDigest Enzymes (EcoRI, KpnI, NotI, XhoI) including buffers, and Rapid DNA Ligation Kit for molecular cloning were purchased from Thermo Fisher Scientific (Rockford, IL, USA). Plasmid pEGFP-N1 (Clontech, Takara Bio USA, Inc, CA, USA) was generously provided by Dr. Gildert Lauter and Dr. Peter Swoboda from the Department for Biosciences and Nutrition, Karolinska Institutet. Mammalian TrxR1 was purchased from IMCO (Stockholm, Sweden) and Sigma-Aldrich (Cas no:9074-14-0, Product number T9698, Darmstadt, Germany).

### Cell culture and growth condition

HEPG2 cells and Hep3B cells were purchased from ATCC (Wesel, Germany) and Huh7 cells were a generous gift from Dr. Camilla Pramfalk (Division of Clinical Chemistry, Karolinska University Hospital Huddinge, Sweden). These cell lines were maintained in EMEM (ATCC) supplemented with 10% heat-inactivated FBS (Gibco, Paisley, United Kingdom), grown under 5 % CO_2_ at 37° C without the addition of antibiotics. Cell lines were regularly tested for mycoplasma infection and kept negative. Cell numbers were assessed using a TC 20^™^ automated cell counter (BioRad, Portland, ME).

Human hepatocytes were isolated as previously described (20) from the surplus surgical waste within the scope of informed consent given by patients undergoing liver resection, and provided by Dr. Ewa Ellis (CLINTEC, Karolinska University Hospital Huddinge, Sweden). The study protocol for the acquisition of cells conformed to the ethical guidelines of the 1975 Declaration of Helsinki and was approved by the Regional Ethics Committee for Human Studies, Stockholm, Sweden, **permit number 2017/269-31** and **2019-04866**. Human hepatocyte viability was assessed by the trypan blue exclusion method. The retrieved cells were seeded on collagen-treated vessels (at a density of 150,000 viable cells per cm^2^) and maintained in Williams’ medium E (Sigma, Darmstadt, Germany) supplemented with 1 μM dexamethasone, 25 mM HEPES, 12 nM Insulin, 2 mM glutamine and 100 U/ml penicillin and 100 μg/ml streptomycin (Thermo Scientific, Rockwood, TN, USA).

### MSC cytotoxicity assessment

To determine MSC cytotoxicity, cells were seeded in 96-well plates (BD Falcon, Durham, NC, USA) at a density of 400 cells/mm^2^ for HEPG2 and Hep3B cells and 300 cells/mm^2^ for Huh7 cells 24 h before MSC treatment. For the determination of the half-maximal inhibitory concentration (IC50), MSC was serially diluted to 10 different concentrations. The ATP content of the lysed cells, which is proportional to the number of viable cells, was measured after 48 and 72 h using the luminescence-based CellTiter-Glo^®^2.0 cell viability assay kit (Promega, Madison, WI, USA) according to the manufacturer’s instructions with an FLx100 Luminometer (CLARIOSTAR^®^, Ortenbery, Germany).

### Cloning of KYAT1 overexpression vectors

To obtain a KYAT1 overexpression vector, the KYAT1 wild-type coding sequence was PCR-amplified from cDNA and inserted into the pEGFP-N1 backbone by XhoI and EcoRI restriction digestion. Subsequently, the human PGK promoter and puromycin resistance gene sequences were amplified from pLKO.1 and inserted into the pEGFP-N1 backbone containing the KYAT1 expression cassette by KpnI and NotI digestion. An empty vector was created by inserting the PCR-amplified Puromycin resistance gene expression cassette into pEGFP-N1 directly. Thus, the eGFP-coding sequence was replaced with the sequence encoding the puromycin resistance gene in both the KYAT1 overexpression vector and the empty control vector.

### Transfection of HCC cell lines

#### KYAT1 overexpression via plasmid

HEPG2 cells were transfected with 2 μg of either KYAT1 overexpression vector or an empty vector using Lipofectamine 3000 (Invitrogen, Camarillo, CA, USA) according to the manufacturer’s protocol. Twenty-four-hour post-transfection, the transfection medium was removed, and the cells were seeded at a density of 400 cells/mm^2^ in 96-well plates for subsequent MSC IC50 titration. Successful overexpression of KYAT1 in transfected HEPG2 cells was verified by RT-PCR and Western Blot.

#### KYAT1 over-expression via formulated (lipid nanoparticle encapsulated) and unformulated KYAT1mRNA

Complete N1-methylpseudouridine substituted mRNA was synthesized as previously described (21). After purification, the mRNA was diluted in citrate buffer to the desired concentration and frozen. mRNA sequences of eGFP and KYAT1 were supplied by Moderna Inc. (Cambridge, MA, USA). KYAT1 mRNA with pseudouridine incorporating a 3XmiR122 antisense site in the same relative locations within the 3’ UTR as described by Ruchi et al (18) was purchased from TriLink Biotechnologies (San Diego, CA, USA). These were designated as either unformulated mRNA or formulated mRNA where the latter constructs were encapsulated within specially designed lipid nanoparticles (LNP) as described previously (22). All formulations were confirmed to be between 80 nm – 100 nm particle size, greater than 80% of RNA encapsulation, and <10 EU/mL endotoxin. For transfection, cells were seeded at a density of 400 cells/mm^2^ or 300 cells/mm^2^ in 60 mm dishes (Sarstedt, North Rhine-Westphalia, Germany) and allowed to grow to 80-90% confluency. Formulated LNP was added at the desired concentration in μg/mL directly to the cell culture media, whereas unformulated mRNA was combined with the respective amount of Lipofectamine 3000 reagent according to the manufacturer’s protocol. Transfected cells were incubated for 4 hours followed by washing with PB S with the subsequent addition of fresh medium. Cells were grown for an additional 20 h before they were harvested for protein isolation. To study the protein expression levels over time, cells were transfected with 0.5 μg/mL LNP particles as described above. Transfected cells were harvested at 24, 48, 72, 96, and 120 h post-transfection. To study protein saturation effects cells were transfected with varying concentrations of KYAT1 ranging from 0.1 to 1μg/mL and harvested at 48 h post-transfection. These cell lysates were subjected to western blot for protein quantification and transamination assay for enzyme activity. For the determination of IC50 value, cells were further treated as described above under the MSC cytotoxicity section.

### Quantitative real-time PCR

Transfected cells were harvested after 24 and 48 h post-transfection. Total RNA was isolated using the RNeasy Plus Mini kit (Qiagen, Hilden, Germany) according to the manufacturer’s instructions. RNA content was measured using a NanoDrop spectrophotometer ND-1000 with subsequent cDNA synthesis using the Ominscript RT kit (Qiagen, Hilden, Germany) according to the provided protocol. RT-PCR was performed using the iQ SYBR green supermix (Bio RAD, Portland, ME). Expression analysis was executed in triplicates and the expression of HPRT was used as a reference gene. PCR primers were as follows:

***KYAT1*** F: CACGCTGTCAGTGGAGACTT, R: TATCTCCTGACCCAGCAGCT,
***HPRT1*** F: GCAGACTTTGCTTTCCTTGG, R: TATCCAACACTTCGTGGGGT,

For microRNA expression analysis, 10ng of RNA was used for synthesizing cDNA using a Taqman advanced microRNA cDNA synthesis kit (Applied Biosystems Inc., Foster City, CA) followed by Taqman advanced microRNA assay (Applied Biosystems Inc., Foster City, CA) and qRT-PCR was performed by TaqMan fast advanced master mix (Applied Biosystems Inc., Foster City, CA). miR122-5p (#477855-mir) expression analysis was executed in triplicate and the expression of miR192-5p (#478262-mir) and miR16-5p (#477860-mir) were used as reference genes. All miRNAs probes were bought from Applied Biosystems Inc., Foster City, CA.

### Quantification assay for total protein content

Samples were lysed in RIPA buffer (Sigma, Darmstadt, Germany) in the presence of 1 mM phenylmethanesulfonyl fluoride (PMSF) and 1% Protease inhibitor cocktail mix (Sigma, Darmstadt, Germany). Further, lysed cells were sonicated at 4 °C for 30 s with 1- to 2-s pulses. The proteins were harvested by centrifugation at 13,000 rpm for 10 min at 4 °C, the supernatant collected, and protein concentration determined by the Pierce™ Bicinchoninic acid (BCA) Protein Assay Kit (Thermo Fischer Scientific, Rockford, IL, USA) according to the manufacturer’s instructions.

### Western blot analysis

Separation of 20 μg or 50 μg protein from cell lysate was performed on a 10% Mini-PROTEAN^®^ TGX^™^ gel (BioRad, Portland, ME) and subsequently transferred onto Immunoblot PVDF 0.45μm membranes (BioRad, Portland, ME) by semi-dry transfer. The membranes were blocked in 5 % milk in 1XTBST and immunoblotted with primary antibodies against KYAT1 (1:1000, anti-CCBL1, #C4622) (Sigma, Darmstadt, Germany) overnight at 4°C. Vinculin (#V284) (1:5000, Millipore, Billerica, MA) was used as a loading control. Membranes were washed and incubated with secondary infrared fluorescent IRDye^®^ antibody (LI-COR^®^, 1:10000) (Nebraska, USA). Subsequently, membranes were washed 3 times with 1XTBST, and blot images were acquired in an Odyssey Fc (LI-COR^®^) (Nebraska, USA) Imaging system. Quantification of the protein levels was determined in the Odyssey Image software (LI-COR^®^) (Nebraska, USA) by normalizing the fluorescence intensity of detected KYAT1 antibody signals to the loading control vinculin signal.

### KYAT1 enzyme activity assay

KYAT1 transamination activity was measured with whole-cell lysate according to the published procedure (23).

Coupled β-elimination activity assay was measured with whole-cell lysate according to the published procedure (24). To assess the inhibitory effects of transamination and β-elimination enzyme activity, the test compounds were added at the desired concentration as indicated with KYAT1 overexpressed whole cell lysate (20 μg).

### Statistical analysis

Results are expressed as mean ± SD and represented in Box and Whisker plots or Violin plots showing median, 25- and 75-percentiles. The analysis was performed by one-way ANOVA with 95% confidential interval followed by Tukey’s or Dunnett’s multiple comparison test (ns= not significant, *p<0.05, **p<0.01, ***p<0.01 & ****p<0.0001) compared to control when specified. Statistical differences between IC50 values were determined by fitting nonlinear regression slopes on independent experiments (n≥3) and subjecting the data to one-way ANOVA followed by Tukey’s multiple comparison test. Data were analyzed with GraphPad Prism software, version 8.3.3 (GraphPad Software Inc, San Diego, CA, USA).

## Results

### Growth inhibitory effect of MSC in HCC cell lines and normal primary hepatocytes

MSC cytotoxicity was evaluated in the three HCC cell lines HEPG2, Hep3B, Huh7 cells and in freshly isolated primary human hepatocytes. Exposure of HCC cell lines and primary human hepatocytes to MSC resulted in IC50 values of 1152 ± 188, 1875 ± 171, 1294 ± 336 and 1218 ± 364 μM, respectively, indicating relatively similar cell sensitivities to MSC exposure in all the cell lines except for Hep3B cells (**Fig. 1A**). The transamination activity was similar in HCC cell lines and primary hepatocytes. The calculated enzyme activity was 0.8±0.3, 0.6±0.2, 1.4±0.3 and 0.8±0.7 nmol of PPA-enol formed/min/mg of protein in HEPG2, Hep3B, Huh7 and primary hepatocytes, respectively (**Fig. 1B**). Western blot analysis confirmed a low endogenous expression level of KYAT1 in HCC cell lines as compared to primary hepatocytes (**Fig. 1C**). The reduced sensitivity (IC50 value) measured in Hep3B correlated with reduced KYAT1 expression levels. Similarly, primary hepatocyte (37-fold increased expression compared to Hep3B cells), as well as HEPG2 and Huh7 (3- and 4-fold increased expression compared to Hep3B cells, respectively) showed elevated expression levels and thus related increased sensitivity to MSC.

**Figure 1.**
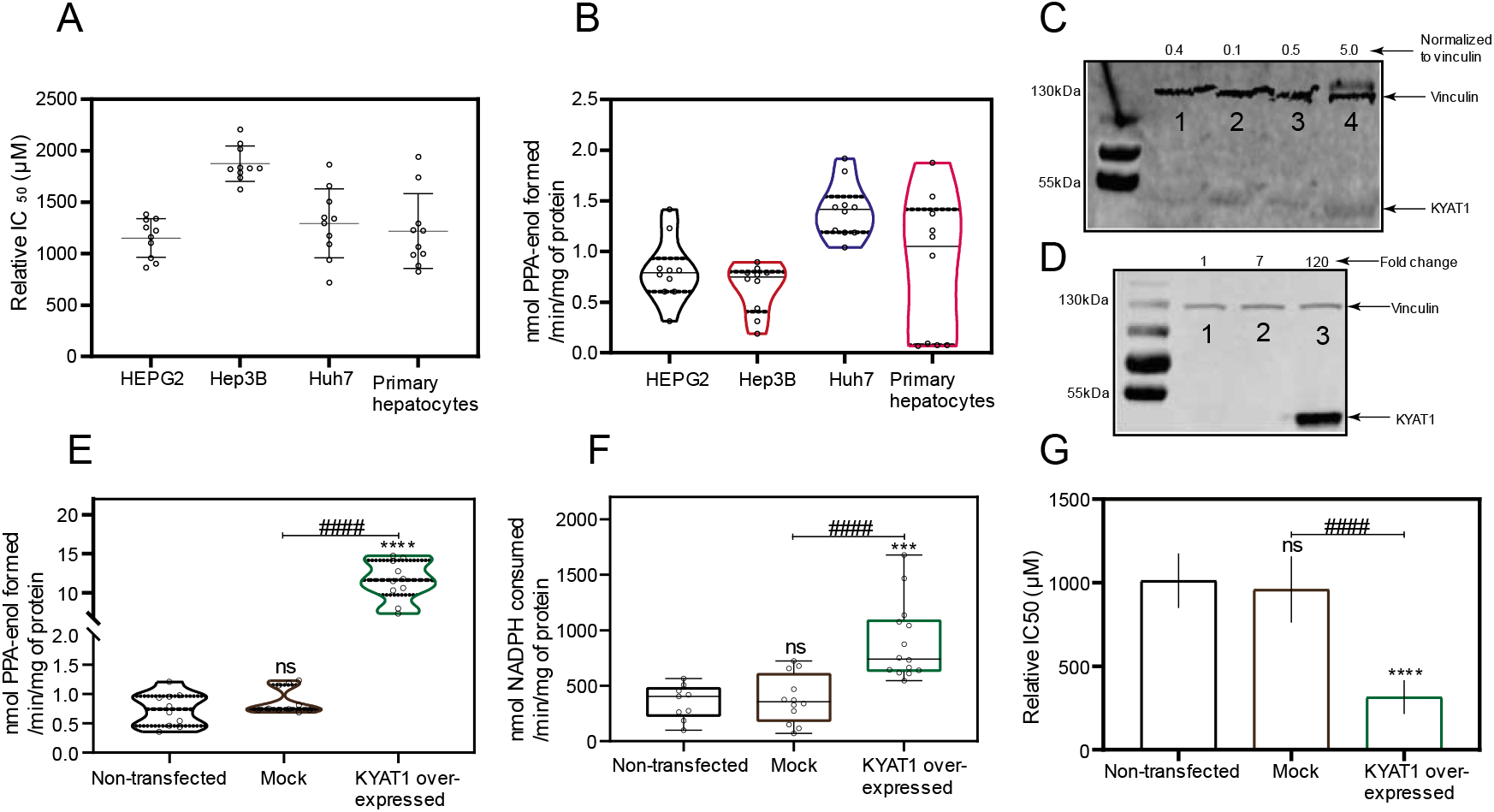
KYAT1 expression and MSC cytotoxicity in cancer and normal human liver hepatocytes. **(A)** MSC cytotoxicity in three different hepatocellular carcinoma cell lines (HEPG2, Hep3B and Huh7) and freshly isolated human primary hepatocytes at 72 h (n=10). **(B)** Transamination activity of KYAT1 in three HCC cell lines and human primary hepatocyte (n=10). **(C)** Western blot showing the endogenous expression of KYAT1 protein in three HCC cell lines and human primary hepatocytes. 1) HEPG2, 2) Hep3B, 3) Huh7 and 4) freshly isolated human primary hepatocytes. **(D)** Over-expressed KYAT1 enzyme was verified by western blot after 48 h of transfection in HEPG2 cells. 1) control, 2) mock-transfected and 3) KYAT1 over-expressing plasmid-transfected cells. **(E)** Transamination activity (n=10) and **(F)** Beta-elimination (n=9-14) activity in control, mock and KYAT1 over-expressed HEPG2 cell lysate. **(G)** KYAT1 over-expression increases the sensitivity of MSC mediated cytotoxicity at 72 h in HEPG2 cells (n=14). **(C) & (D)** KYAT1 48kDa and Vinculin 124kDa were used as a loading control and 20 μg of protein from cell lysate was used. **(E-G)** Graph represent mean ± SD, statistical analysis performed with one-way ANOVA with 95% confidential interval followed by Tukey’s multiple comparison test (ns = not significant, \**p* < 0.05, \*\**p* < 0.01, \*\*\**p* < 0.001 and \*\*\*\**p* < 0.0001 compared with untransfected control and # p<0.05, ## p<0.01, ### p<0.001 & #### p<0.0001 compared with mock).

### KYAT1 plays a major role in MSC metabolism in HCC cell lines

KYAT1 was overexpressed in HEPG2 cells using a plasmid that encoded the full-length enzyme. After 48 h post-transfection, overexpression was confirmed by western blot (**Fig. 1D**). Transamination (**Fig. 1E)** and β-elimination (**Fig. 1F**) activities were quantified in KYAT1 over-expressed cell lysate. Transamination activity (0.7±0.3, 0.9±0.2 and 11.6±2.6 nmol of PPA-enol formed/min/mg of protein in un-transfected, mock and KYAT1 over-expressed cells, respectively) and β-elimination activity (0.4±0.2, 0.4±0.2 and 0.9±0.3 μmol of NADPH consumed/min/mg of protein in un-transfected, mock and KYAT1 over-expressed cells, respectively) were significantly increased in KYAT1 over-expressed HEPG2 cells. KYAT1 overexpression sensitized HEPG2 cells to MSC treatment, resulting in significantly reduced IC50 values (316±102 μM) as compared to non-transfected and mock-transfected controls (1013±163 and 960±198, respectively) (**Fig. 1G**). In contrast, siRNA-mediated knockdown provided a protective effect on MSC-mediated cytotoxicity in Huh7 cells (Supplementary Fig. S1A & 1B). Overall, our results demonstrate that KYAT1 over-expressing cells displayed significantly higher transamination and β-elimination activities, combined with significantly increased cytotoxicity upon exposure to MSC.

### Intracellular delivery of KYAT1mRNA using lipid nanoparticles

In addition to the plasmid (DNA)-based over-expression system, we investigated the delivery of mRNA encoding for KYAT1, encapsulated in lipid nanoparticles (LNPs). HEPG2 cells were exposed to increasing concentrations of KYAT1-encoding mRNA-LNPs. The normalized signal intensity of western blot revealed an increase in KYAT1 protein expression from the lowest concentration of 0.1 μg/mL (60-fold compared to control) to 1.0 μg/mL sample (100-fold compared to control) (**Fig. 2A**). Protein determined over time in HEPG2 cells transfected with 0.5 μg/mL of KYAT1 mRNA, revealed 190-fold increased KYAT1 expression levels at 48 h compared to control, with a small and slow decline over time (**Fig. 2B**). Enzyme activity assays correlated with the protein expression level (**Fig. 2C & 2D, Supplementary Table S1A & S1B**), showing minimal differences in transamination activity in HEPG2 cells transfected with varying KYAT1mRNA concentrations at a fixed time point, and fixed concentration with varying time points. As a consequence of the proven over-expression of KYAT1 following efficient LNP mRNA delivery, we decided to limit our further exposed concentration to 0.2 and 0.5 μg/mL of mRNA.

**Figure 2.**
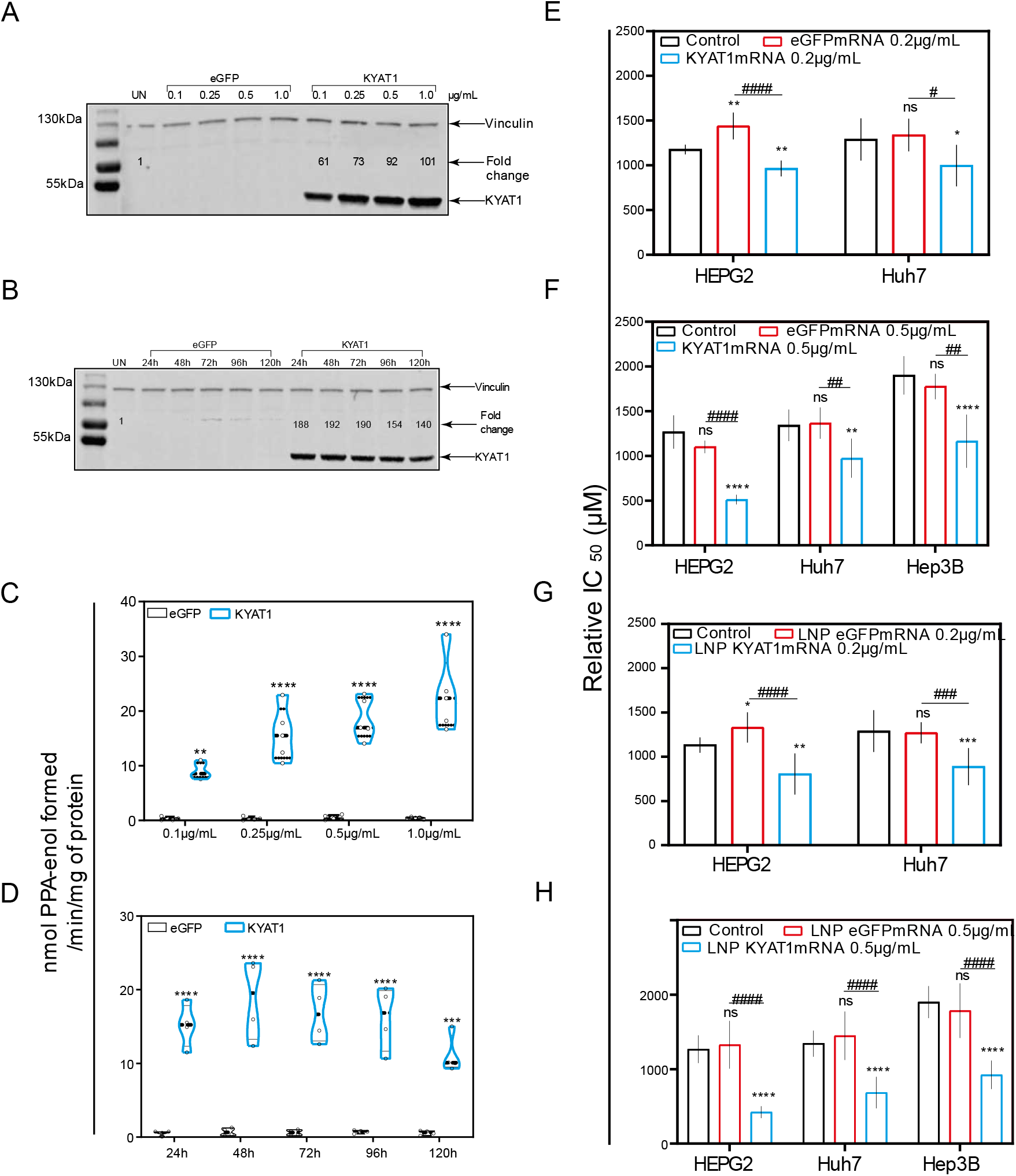
KYAT1 upregulation increases the efficacy of MSC cytotoxicity in HCC cell lines. Westernblot showing the expression of KYAT1 in HEPG2 cells transfected with **(A)** varying concentrations (0.1,0.25, 0.5 and 1 μg/mL) of KYAT1mRNA or eGFPmRNA encapsulated in LNP at 48h. **(B)** 0.5 μg/mL of LNP encapsuled KYAT1mRNA or eGFPmRNA at different time points (24, 48, 72, 96 and 120 h). **(C) T**ransamination activity of KYAT1 was determined for samples at varying concentration at fixed time (n=5) and **(D)** Fixed concentration (0.5 μg/mL) at varying time (n=4). **Dose-dependent effect of MSC at 72 h in HCC cell lines transfected with (E)** mRNA encoding KYAT1 or eGFP (0.2 μg/mL), (n=6-11) **(F)** mRNA encoding KYAT1 or eGFP (0.5 μg/mL) (n=5-11), **(G)** LNP encapsulated KYAT1 or eGFP mRNA (0.2 μg/mL) (n=6-11) & **(H)** LNP encapsulated KYAT1 or eGFP mRNA (0.5 μg/mL) (n=6-11). **(A)** & **(C)** KYAT1 48KDa and Vinculin 124KDa used as loading control, UN = un-transfected control. **(E-H)** eGFPmRNA and non-transfected cells were used as a control with respective concentrations as mentioned above. Graph represent mean ± SD, statistical analysis performed with **(B) (D)** Unpaired t-test & **(E-H)** one-way ANOVA with 95% confidential interval followed by Tukey’s multiple comparison test (ns = not significant, * *p* < 0.05, \*\**p* < 0.01, *** *p* < 0.001 and \*\*\*\**p* < 0.0001 compared with control and # p<0.05, ## p<0.01, ### p<0.001 & #### p<0.0001 compared with eGFPmRNA).

### Increased MSC cytotoxicity in HCC cell lines upon KYAT1mRNA delivery using lipid nanoparticles or lipofectamine

MSC cytotoxicity was measured in all three HCC cell lines following transfection with KYAT1 mRNA. Both mRNA concentrations (0.2 and 0.5 μg/mL) resulted in enhanced sensitivity to MSC cytotoxicity in all three HCC cell lines **(Fig. 2E–2H)**. Unformulated KYAT1mRNA (0.2μg/mL) sensitized HEPG2, Hep3B and Huh7 cells to MSC significantly as compared to untransfected or eGFPmRNA transfected cells (**Fig. 2E, Table 1A**). As hypothesized, increased KYAT1mRNA concentrations (0.5μg/mL) further sensitized HEPG2, Hep3B and Huh7 cells to MSC treatment (**Fig. 2F, Table 1A**). Furthermore, cells transfected with the LNP-encapsulated KYAT1 showed a higher or similar sensitization effect towards MSC in the presence of 0.2 and 0.5 μg/mL of KYAT1mRNA (**Fig. 2G & 2H & Table 1B**). Cells transfected with LNP-based mRNA delivery showed higher sensitivity towards MSC cytotoxicity as compared to lipofectamine-transfected cells (**Table 1A & Table 1B**). Our experimental data also showed that LNP delivery *per se* did not affect MSC uptake and cytotoxicity in HCC cell lines (**Supplementary Fig. S1C**).

**Table 1A.** Relative IC_50_ values of HEPG2, Huh7 and Hep3B cells transfected with 0.2 μg/mL and 0.5 μg/mL of eGFPmRNA- or KYAT1mRNA using lipofectamine 3000. Results are presented as mean ± SD from triplicate measurements from at least five independent experiments. Resultant IC_50_ values are denoted in μM.

|  | Cells transfected via lipofectamine3000 |  |  |  |  |
| --- | --- | --- | --- | --- | --- |
| HCC cell lines | Untransfected cells | eGFPmRNA control 0.2 µg/mL | KYAT1mRNA control 0.2 µg/mL | eGFPmRNA control 0.5 µg/mL | KYAT1mRNA control 0.5 µg/mL |
| HEPG2 | 1200±157 | 1440±151 | 964±88 | 1101±71 | 511±55 |
| Huh7 | 1315±204 | 1339±183 | 997±232 | 1366±176 | 973±219 |
| Hep3B | 1900±216 | ND | ND | 1776±142 | 1164±297 |

ND = not determined

**Table 1B.** Relative IC_50_ values of HEPG2, Huh7 and Hep3B cells transfected with 0.2 μg/mL and 0.5 μg/mL of LNP encapsulated eGFPmRNA- or KYAT1mRNA. Results are presented as mean ± SD from triplicate measurements from at least five independent experiments. Resultant IC _50_ values are denoted in μM.

|  | Cells transfected via lipid nanoparticles (LNP) |  |  |  |  |
| --- | --- | --- | --- | --- | --- |
| HCC cell lines | Untransfected cells | eGFPmRNA 0.2 µg/mL | KYAT1mRNA 0.2 µg/mL | eGFPmRNA 0.5 µg/mL | KYAT1mRNA 0.5 µg/mL |
| HEPG2 | 1200±157 | 1330±171 | 804±232 | 1327±321 | 422±80 |
| Huh7 | 1315±204 | 1270±120 | 888±211 | 1447±326 | 644±216 |
| Hep3B | 1900±216 | ND | ND | 1883±330 | 924±191 |

ND = not determined

### Pharmacological modulation of KYAT1 activity

To modulate the β-elimination or transamination activity, several pharmacological modulating compounds were tested and four different KYAT1 inducers and inhibitors were selected. Transamination and β-elimination activity assays were used to quantify KYAT1 activity in the presence of selected inducers and inhibitors in LNP transfected cells (**Supplementary Fig. S2A-2E**). HEPG2, Hep3B and Huh7 cells were transfected with 0.5 μg/mL eGFP or KYAT1mRNA encapsulated in LNP. The day after, transfected cells were exposed to MSC alone or in combination with inducers/inhibitors for an additional 72 hrs. Our results revealed that the KYAT1 inhibitor AOAA reversed MSC induced cytotoxicity in all the cell lines and that 85% of cells persisted viable at 2 mM of MSC (**Fig. 3A–3C**). In contrast, MSC combined with the α-ketoacid analog PPA (phenylpyruvic acid) significantly increased the cytotoxic effects in KYAT1 over-expressing cells, as compared to MSC alone in all three HCC cell lines (**Fig. 3A–3C**). Furthermore, the combination of MSC with IPA (indole-3-pyruvic acid) resulted in significant cytotoxic effects in KYAT1 over-expressing Huh7 and Hep3B cells compared with MSC alone (**Fig. 3B & 3C**). Finally, KMB (α-Keto-γ-(methylthio) butyric acid) in combination with MSC did not yield any significant additive effect (**Fig. 3A–3C, Supplementary Table S2**). KYAT1 inducers and inhibitors did not significantly alter MSC cytotoxicity in primary hepatocytes, except AOAA which protected MSC induced cytotoxicity (**Supplementary Fig. S2F**). In summary, the addition of α-ketoacid analogs in combination with MSC resulted in significantly increased cytotoxic effects only in HCC cell lines.

**Figure 3.**
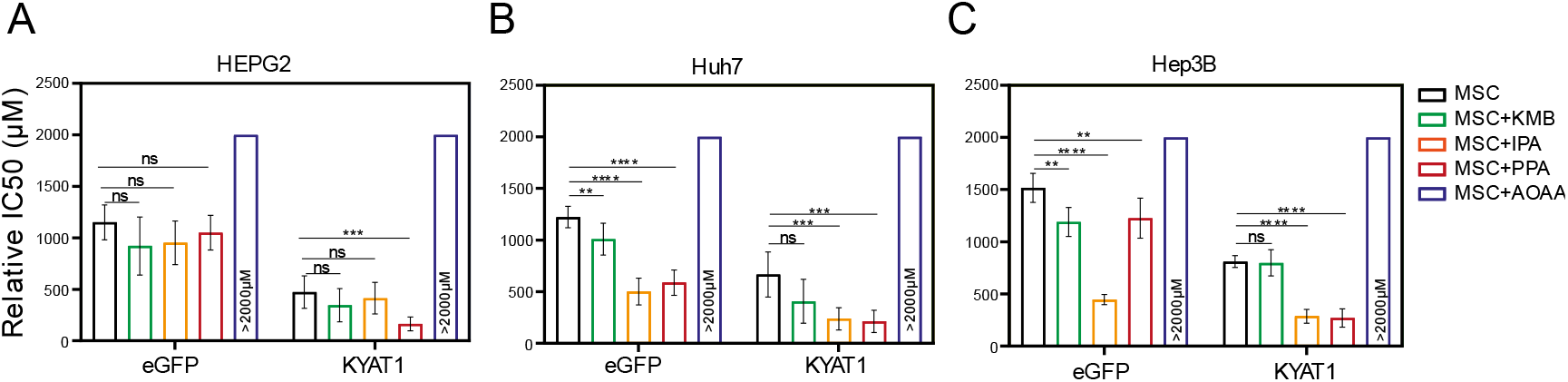
LNP mediated KYAT1 over-expression in combination with α-ketoacids increased the efficacy of MSC treatment in HCC cell lines. Efficacy of MSC co-incubating with α-ketoacids such as phenylpyruvic acid (PPA) (400μM), indole-3-pyruvic acid (IPA) (200μM), α-Keto-γ-(methylthio) butyric acid (KMB) (100μM) and KYAT1 inhibitor aminooxy acetic acid (AOAA) (1mM) in cells transfected with KYAT1mRNA encapsulated in LNP (0.5 μg/mL) in three different HCC cell lines at 72 h (A) HEPG2 (n=8-10), (B) Hep3B (n=6-8) & (C) Huh7 (n=6-8). (A-C) LNPeGFPmRNA transfected cells were used as a control with respective concentrations as above. Graph represent mean ± SD, statistical analysis performed with one-way ANOVA with 95% confidential interval followed by Tukey’s multiple comparison test (ns = not significant, * *p* < 0.05, \*\**p* < 0.01, \*\*\**p* < 0.001 and \*\*\*\**p* < 0.0001 compared with control).

### Antisense microRNA restricts MSC induced cytotoxicity to tumor cells

To restrict over-expression of KYAT1 to only tumor cells, antisense sequences of specific microRNAs commonly downregulated in tumors were added in the 3’end of the coding region of KYAT1. Since selected microRNAs are tissue and disease-specific, we focused our attention on microRNA122 (miR122). A schematic representation of how miR122 may help us to target only HCC is presented in **Fig. 4A**. Endogenous miR122 expression levels were initially quantified in human hepatocytes isolated from healthy donors and HCC, and compared with HEPG2, Hep3B, and Huh7 cell lines (**Fig. 4B, 4C & 4D)**. The calculated Ct values for HEPG2, Hep3B and Huh7 cells were 32.7±0.2, 29.4±0.3 and 19.6±0.1, and the corresponding fold changes were 1, 6.5±2.5 and 4496±887, respectively (**Fig. 4B)**. When miR122 level of expression was assessed in neoplastic tissue and compared with surrounding “normal” parenchyma, we measured 280000-, 659- and 81-fold increased miR122 expression in normal liver hepatocytes compared to neoplastic tissue in three different tested donors (**Fig. 4C)**. The calculated Ct value for freshly isolated human primary HCC (n=5) and primary hepatocyte (normal) cells (n=12) were 28.4±3.7 and 21.4±2.6, respectively, *i.e*. primary hepatocytes displayed a 17-fold higher miRNA122 expression compared to primary HCC (relative expression of HCC and normal hepatocytes were 5024±6882 and 84554±101817 respectively with p-value 0.02) (**Fig. 4C)**. Huh7 cells were characterized by elevated levels of miR122 expression compared to Hep3B (688-fold higher in Huh7) and HEPG2 (4496-fold higher in Huh7) cells. In this respect, Huh7 cell lines showed a comparable expression pattern to normal hepatocytes and thus we considered that they may serve as a relative control for normal hepatocytes. As a proof-of-concept, we delivered mRNA encoding for KYAT1 with three additional microRNA122 target sites (3X122 KYAT1mRNA) into HEPG2, Hep3B, Huh7 and freshly isolated primary hepatocytes. After 48h, KYAT1 transamination activity was significantly reduced in Huh7 cells (**Fig. 4E)** and freshly isolated primary hepatocytes (**Fig. 4F)** transfected with 3X122 KYAT1mRNA as compared to KYAT1mRNA transfected cells (**Fig. 4E & 4F)**. Indeed, Huh7 cells transfected with 3X122 KYAT1mRNA displayed a 70% reduction in protein level (**Fig. 4G & 4H)**. In contrast, Hep3B and HEPG2 cell lines did not display any significant reduction in KYAT1 activity (**Fig. 4G & 4H)**.

**Figure 4.**
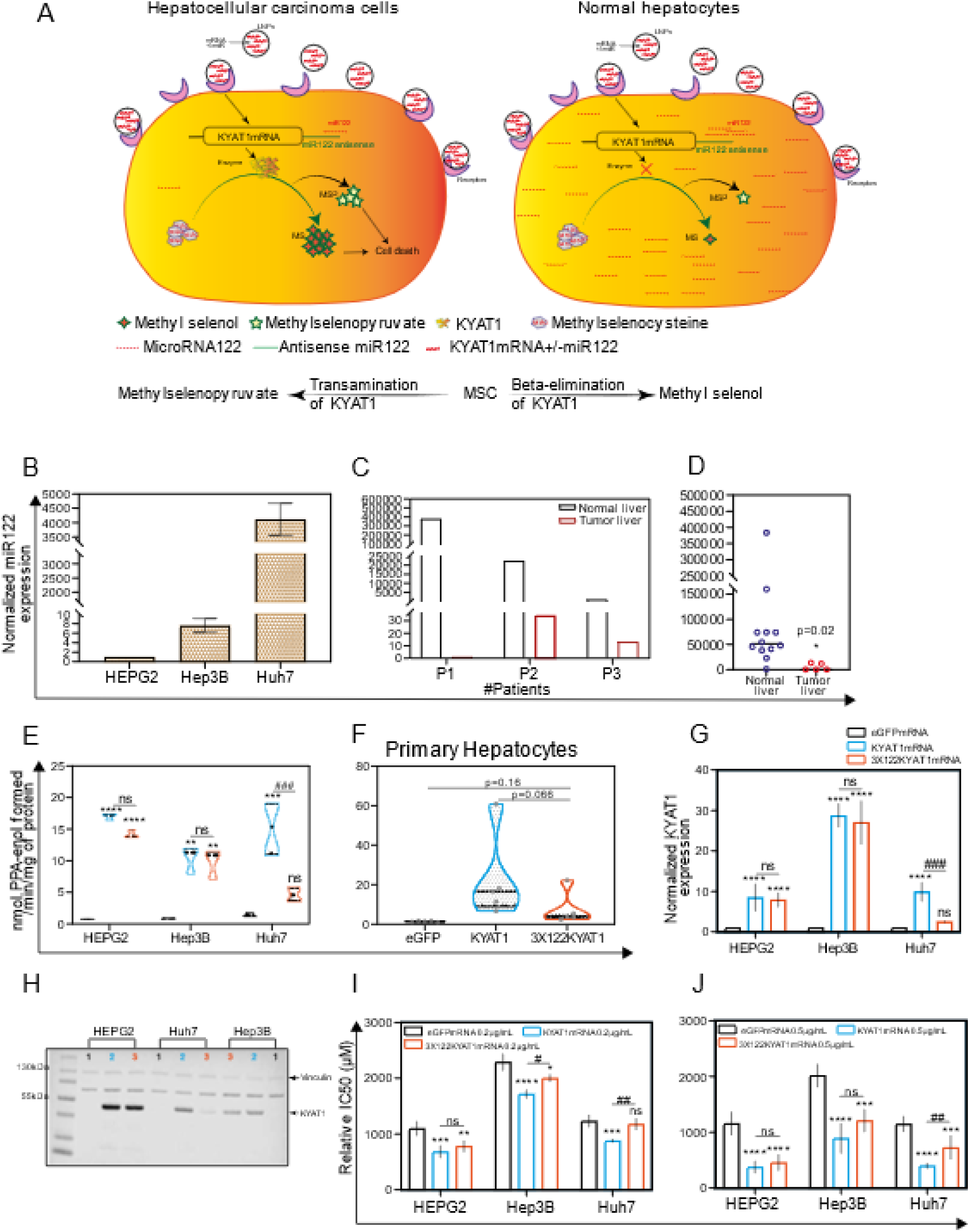
Endogenous expression of miRNA act as a targeted protein suppressor in cells transfected with mRNAs tagged with antisense miRNA target site. (A) Schematic representation of MicroRNA (miR122) assisted MSC metabolism via KYAT1 enzyme in cancer and normal hepatocytes. (B) Endogenous miR122 expression in HEPG2, Hep3B and Huh7 cell lines by qRT-PCR (n=4). (C) miR122 expression in human primary HCC compared with surrounding “normal” primary −hepatocytes from three different patients (n=3). (D) Endogenous miR122 expression in freshly isolated healthy human hepatocytes (n=12) and human primary HCC (n=5). (E) Transamination activity of KYAT1 in cells transfected with eGFP, KYAT1 and 3XmiR122 KYAT1 over-expressing mRNA in HEPG2, Hep3B and Huh7 cells (n=3-4). (F) Transamination activity of KYAT1 in freshly isolated human hepatocytes transfected with eGFP, KYAT1 and 3XmiR122 KYAT1 over-expressing mRNA (n=5). (G) & (H) Western blot showing the expression and quantification of KYAT1 in HEPG2 (n=3), Hep3B (n=3) and Huh7 (n=4) cells 48 h after transfected with KYAT1mRNA and 3XmiR122 KYAT1mRNA. (I) & (J) MSC cytotoxicity in three different HCC cell lines (HEPG2, Hep3B & Huh7) transfected with KYAT1 with or without incorporation of antisense miR122 target site. (I) 0.2 μg/mL & (J) 0.5 μg/mL of mRNA encoding eGFP, KYAT1 & 3XmiR122 KYAT1. (B) (C) & (D) Since the HEPG2 cells have no or minimal expression of miR122, we used HEPG2 as a reference to calculate the miR122 expression in fold changes. Expression was normalized with miR192 and miR16. (H) KYAT1 48KDa and Vinculin 124KDa used as a loading control, 1) eGFP, 2) KYAT1 & 3) 3XmiR122 KYAT1. (I) & (J) eGFPmRNA was used as a control with respective concentrations as above (n=5-7). Graph represent mean ± SD, statistical analysis performed with (C) Unpaired t-test with Welch’s correction (E) (F) (H) (I) & (J) one-way ANOVA with 95% confidential interval followed by Tukey’s multiple comparison test (ns = not significant, * *p* < 0.05, \*\**p* < 0.01, *** *p* < 0.001 and \*\*\*\**p* < 0.0001 compared with eGFP and # p<0.05, ## p<0.01, ### p<0.001 & #### p<0.0001 compared with KYAT1).

HEPG2, Hep3B and Huh7 cell lines were transfected with two different concentrations of KYAT1mRNA (0.2 and 0.5 μg/mL), with or without 3XmiR122 target sites. The relative IC50 values of MSC were measured at 72h. A mild sensitization at 0.2 μg/mL of KYAT1 induction and significant sensitization at 0.5 μg/mL was observed with MSC in all three HCC cell lines (**Fig. 4I & 4J)**. Cells transfected with 0.2 μg/mL 3X122 KYAT1mRNA in Hep3B and Huh7 cells showed a significant protective effect against MSC cytotoxicity, which was supported by qRT-PCR analysis (see miR122 expression in Hep3B cells) (**Fig. 4B)**. At 0.2 μg/mL and 0.5 μg/mL concentration, significant resistance towards MSC cytotoxicity was observed, limited to Huh7 cells (**Fig. 4I & 4J)**. The calculated relative IC50 values at 0.2 μg/mL and 0.5 μg/mL of eGFPmRNA, KYAT1mRNA and 3X122KYAT1mRNA transfected cells are presented in **Table 2**. Altogether, our results indicate that miRNA antisense targets are efficient in achieving tumor-specific cytotoxicity in cell types where miR122 is modulated.

**Table 2.**
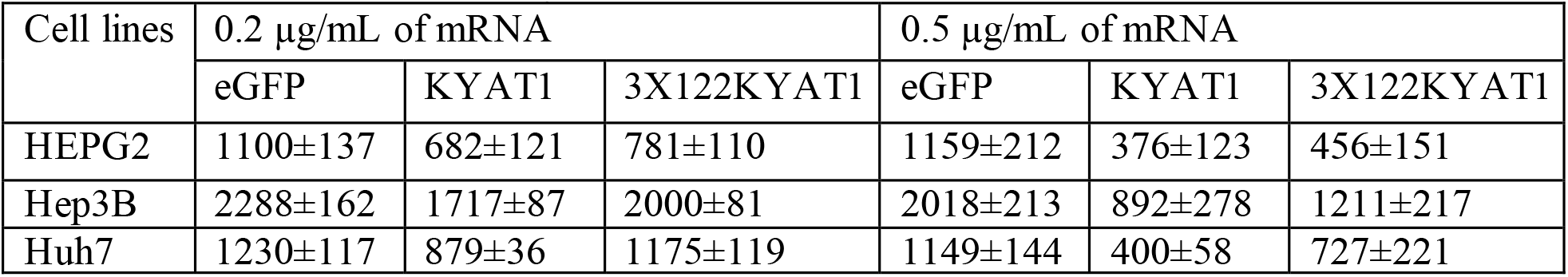
Relative IC_50_ values for MSC at 72 h in HEPG2, Hep3B and Huh7 cells transfected with 0.2 μg/mL and 0.5 μg/mL of eGFPmRNA, KYAT1mRNA, and 3X122KYAT1mRNA. Results are represented as mean ± SD. from triplicate measurements from at least five independent experiments. Resultant IC_50_ values are denoted in μM.

### Endogenous miRNA can protect cells from MSC-induced cytotoxicity even in combination with KYAT1 inducers/inhibitors

HCC cell lines were transfected with two different concentrations of KYAT1mRNA (0.2 μg/mL and 0.5 μg/mL) with or without 3XmiR122 target sites. HEPG2 cells transfected with KYAT1 containing 3XmiR122 did not show any protective effect with MSC when co-incubated with α-ketoacid (**Fig. 5A & 5D)**. Hep3B cells at low KYAT1 concentration (0.2 μg/mL) containing 3XmiR122 showed a significant protective effect in combined treatment with IPA and KMB, but these effects were diminished at the higher concentration 0.5μg/mL (**Fig. 5B & 5E)**. Huh7 cells showed significant protective effects at 0.5 μg/mL KYAT1 concentration in combination with IPA and KMB. Interestingly we observed no significant differences at 0.2 μg/mL of KYAT1 transfected cells (**Fig. 5C & 5F)**. Neither Hep3B nor Huh7 cells did show any significant protective effect when exposed to PPA. The calculated relative IC50 values are presented in **Table 3A, 3B & 3C**. Altogether, our data indicate that the off-target expression of KYAT1 protein can be controlled by endogenous microRNA in the presence of KYAT1 inducers/inhibitors except for PPA.

**Figure 5.**
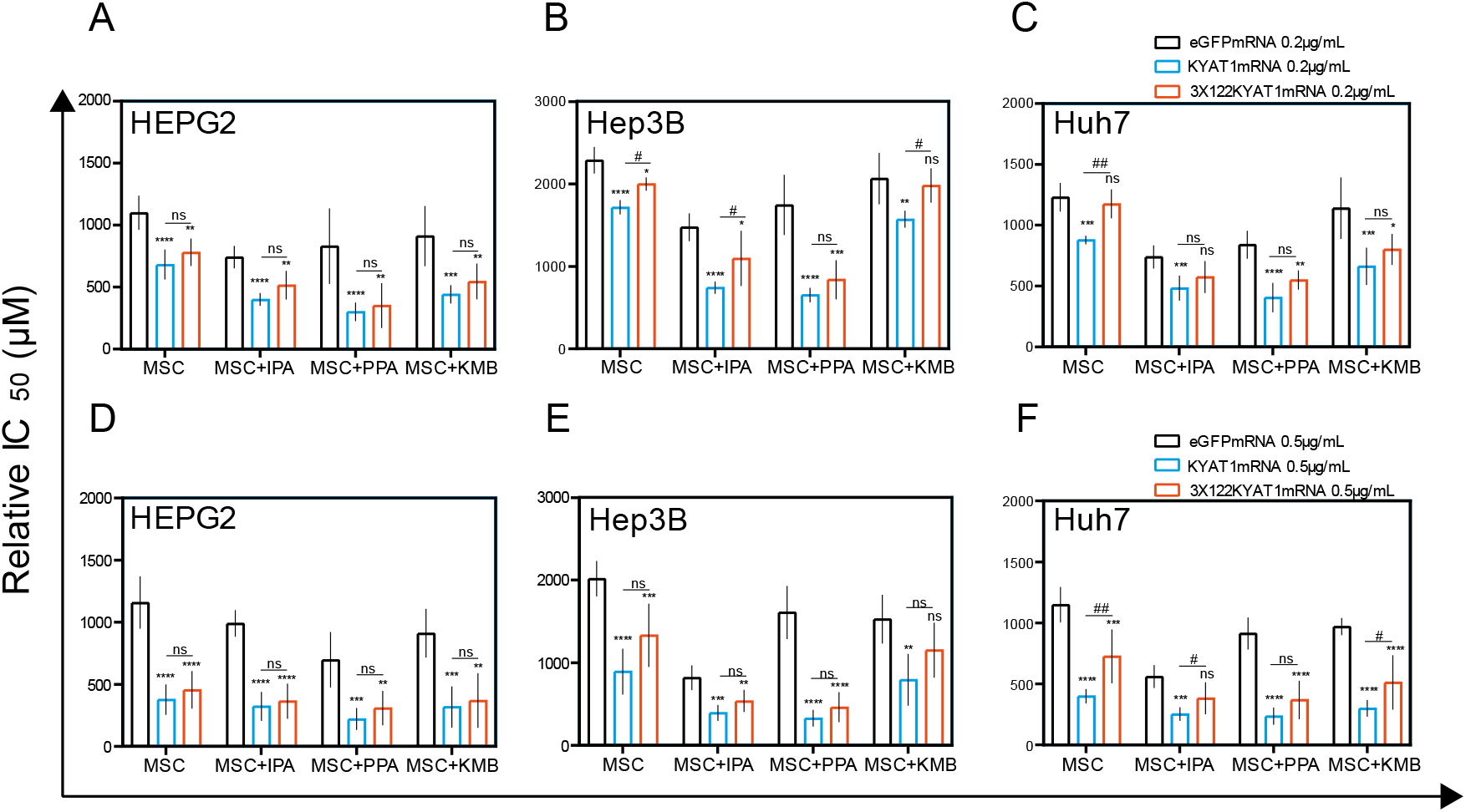
Antisense miR122 target site fusion with KYAT1 alleviates KYAT1 modifiers efficacy with MSC in KYAT1 overexpressed HCC cells. Effect of MSC co-incubated with modifiers in cells transfected with eGFPmRNA, KYAT1mRNA and 3X122KYAT1mRNA. (A) & (D) HEPG2 cells (n=5-7) (B) & (E) Hep3B cells (n=5) and (C) & (F) Huh7 cells (n=5-7). (A) (B) & (C) 0.2 μg/mL and, (D) (E) & (F) 0.5 μg/mL of mRNA encoding eGFPmRNA, KYAT1mRNA & 3X122KYAT1mRNA. Modifiers such as phenylpyruvic acid (PPA) (400μM), indole-3-pyruvic acid (IPA) (200μM) and α-Keto-γ-(methylthio) butyric acid (KMB) (100μM) were co-incubated with MSC and relative IC50 values were calculated at 72 h. Graph represent mean ± SD, statistical analysis performed with one-way ANOVA with 95% confidential interval followed by Tukey’s multiple comparison test (ns = not significant, * *p* < 0.05, ** *p* < 0.01, *** *p* < 0.001 and **** *p* < 0.0001 compared with eGFP and # p<0.05, ## p<0.01, ### p<0.001 & #### p<0.0001 compared with KYAT1).

**Table 3A.** Relative IC_50_ values for MSC alone or in combination with modifiers at 72 h in HEPG2 cells transfected with 0.2 μg/mL and 0.5 μg/mL of eGFPmRNA, KYAT1mRNA, and 3X122KYAT1mRNA. Results are represented as mean ± SD. from triplicate measurements from at least five independent experiments Resultant IC_50_ values are denoted in μM.

| Treatments | HEPG2 Relative IC50 in µM |  |  |  |  |  |
| --- | --- | --- | --- | --- | --- | --- |
|  | 0.2 µg/mL of mRNA |  |  | 0.5 µg/mL of mRNA |  |  |
|  | eGFP | KYAT1 | 3X122KYAT1 | eGFP | KYAT1 | 3X122KYAT1 |
| MSC | 1100±137 | 682±121 | 781±110 | 1159±137 | 376±123 | 456±151 |
| MSC+IPA | 743±89 | 401±50 | 515±116 | 990±107 | 322±116 | 364±141 |
| MSC+PPA | 830±305 | 301±74 | 352±180 | 697±223 | 220±88 | 309±138 |
| MSC+KMB | 912±243 | 442±74 | 546±144 | 911±197 | 318±165 | 369±220 |

**Table 3B.** Relative IC_50_ values for MSC alone or in combination with modifiers at 72 h in Hep3B cells transfected with 0.2 μg/mL and 0.5 μg/mL of eGFPmRNA, KYAT1mRNA, and 3X122KYAT1mRNA. Results are represented as mean ± SD. from triplicate measurements from at least five independent experiments. Resultant IC_50_ values are denoted in μM.

| Treatments | Hep3B Relative IC50 in µM |  |  |  |  |  |
| --- | --- | --- | --- | --- | --- | --- |
|  | 0.2 µg/mL of mRNA |  |  | 0.5 µg/mL of mRNA |  |  |
|  | eGFP | KYAT1 | 3X122KYAT1 | eGFP | KYAT1 | 3X122KYAT1 |
| MSC | 2288±162 | 1717±87 | 2000±81 | 2018±213 | 892±278 | 1331±384 |
| MSC+IPA | 1476±169 | 743±74 | 1097±335 | 818±150 | 392±96 | 537±133 |
| MSC+PPA | 1745±365 | 652±88 | 839±237 | 1609±321 | 328±102 | 461±181 |
| MSC+KMB | 2065±312 | 1571±102 | 1982±93 | 1528±296 | 793±313 | 1152±333 |

**Table 3C.** Relative IC_50_ values for MSC alone or in combination with modifiers at 72 h in Huh7 cells transfected with 0.2 μg/mL and 0.5 μg/mL of eGFPmRNA, KYAT1mRNA, and 3X122KYAT1mRNA. Results are represented as mean ± SD. from triplicate measurements from at least five independent experiments. Resultant IC_50_ values are denoted in μM.

| Treatments | Huh7 Relative IC50 in µM |  |  |  |  |  |
| --- | --- | --- | --- | --- | --- | --- |
|  | 0.2 µg/mL of mRNA |  |  | 0.5 µg/mL of mRNA |  |  |
|  | eGFP | KYAT1 | 3X122KYAT1 | eGFP | KYAT1 | 3X122KYAT1 |
| MSC | 1230±118 | 879±36 | 1175±119 | 1149±144 | 400±58 | 727±221 |
| MSC+IPA | 741±95 | 483±103 | 575±132 | 560±93 | 254±56 | 382±131 |
| MSC+PPA | 840±114 | 406±121 | 550±78 | 914±131 | 236±70 | 387±158 |
| MSC+KMB | 1141±253 | 663±153 | 802±128 | 969±70 | 300±68 | 514±223 |

## Discussion

Since Se-methylselenocysteine (MSC) is a highly interesting prodrug with very favorable pharmacokinetic properties and high per-oral bioavailability, the development of MSC-based strategies for the treatment of cancer is highly relevant. Selenium-based chemotherapeutic regimens are not yet in clinical practice due to the lack of systematic clinical trials and regimens that can provide a high degree of tumor specificity thereby limiting off-target effects. We present herein a novel strategy to achieve efficient and specific tumoricidal effects by combining the prodrug MSC with tumor-specific microRNA-guided overexpression of KYAT1, a strategy that we denote Targeted Enzyme Assisted Chemotherapy (TEAC) (**Fig. 4A)**.

To study the endogenous KYAT1 activity we determined the half-maximal inhibitory concentration (IC50) in hepatocellular carcinoma cell lines (HCC) in comparison to normal human hepatocytes, to establish a feasible study system of diverse responses to MSC treatment. Hep3B cells showed high resistance towards MSC cytotoxicity while HEPG2, Huh7 and normal primary cells showed similar IC50 values. Such differences in sensitivity to MSC treatment could be potentially explained by related endogenous levels of KYAT1 (25). To our knowledge, there have hitherto been limited reports on the roles of KYAT1 in cancer. KYAT1 plays an important role in tryptophan metabolism by metabolizing kynurenine to kynurenic acid (KYNA). KYNA has anti-proliferative activity attributed to its ability to inhibit p21, leading to cell cycle arrest (26). Furthermore, KYNA exhibited clear anti-tumoral effects in human renal cell adenocarcinoma as well as in colon cancer cell models (27, 28).

Overexpression of KYAT1 in HCC cell lines resulted in a significant alteration in the dose-response curve of MSC with a 65-70 % decrease in IC50 values compared to empty vector or un-transfected cells. Our results confirm the preliminary analysis of the KYAT1 −dependent cleavage of MSC (29). To verify the increased enzymatic activity following KYAT1 over-expression, transamination and β-elimination activities were measured in cell lysates. Our results demonstrated a 15-fold higher transamination activity and a 2-fold higher β-elimination activity in KYAT1-overexpressed cell lysates. The sparse β-elimination activity is likely because KYAT1 favors transamination over β-elimination activity (30).

The use of mRNA is preferred over DNA to ectopically overexpress a protein since mRNA will only result in transient, controllable overexpression without risk of incorporation into the genome of the target cell, thus minimizing effects from transgene expression. Currently, RNA-based therapeutics are emerging in clinical studies in treating various diseases including cancer and various genetic disorders (31). We employed an LNP-based delivery system to deliver KYAT1mRNA (32). Our results confirmed that (i) the LNP delivery system was an adequate and efficient tool for successful transfection, (ii) KYAT1 protein levels remained overexpressed for the duration of 120 h and (iii) there was no interference of LNP particles with MSC cytotoxicity.

Upon inducing KYAT1 via mRNA with (formulated mRNA) or without (unformulated mRNA) LNP encapsulation, the transfected HCC cell lines exhibited significantly increased sensitivity towards MSC compared to eGFPmRNA transfected control. The KYAT1 activity depends on several factors, including the nucleophilicity of a substrate, PLP-PMP (co-factor) conversion rate and the presence of α-ketoacid (33, 34). The highly nucleophilic selenium moiety favors β-elimination over transamination which explains why MSC is a preferred substrate for KYAT1 compared to sulphur analogs (29). To explore the impact of α-ketoacid analogs on the efficacy of KYAT-mediated MSC activation, several α-ketoacids were screened. We supplemented the culture medium with indole-3-pyruvic acid (IPA) (35), phenylpyruvic acid (PPA) (36) and α-keto-γ-(methylthio)butyric acid sodium salt (KMB) (30) in the presence of MSC. The effect of IPA was unexpected given that the presence of this analog increased MSC cytotoxicity even without enzyme induction, indicating that the endogenous enzyme was activated by IPA. Han *et al* showed that L-tryptophan and IPA were potent competitive endogenous inhibitors of KYAT1 (37). They further investigated the possible mechanisms underlying the inhibition of IPA using indole-3-acetic acid (IAA) due to structural similarities between these two compounds and reported that IAA was not a substrate for KYAT1. It has been previously demonstrated that the moieties important for the binding within the active site were the indole ring and the carboxyl group (37). This observation strongly suggests that MSC might be able to dock within the active site at the same time as IPA. Thus, the α-ketoacid analog in question may only inhibit the transaminase activity of KYAT1, promoting the production of cytotoxic MS. Unfortunately, IPA could not be used in the β-elimination activity assay because the color of this compound interfered with the β-elimination activity assay and showed a higher absorbance. On the other-hand, KYAT1 activity was significantly increased in the coupled-β-elimination activity assay in the presence of PPA and this explains the enhanced cytotoxicity with MSC (29). Based on these results we conclude that sensitization is due to the self-renewal of PPA in cells. PPA reacts with L-tryptophan to generate L-phenylalanine which in turn reacts with 2-oxobutanoate to regenerate PPA. This continuous supply of PPA could thus represent an efficient pathway to continuously supply PLP required for the β-elimination reaction to proceed. It has previously been reported that the addition of phenylpyruvate to the transamination reaction abrogates the transamination activity of KYAT1 (30).

β-lyase inhibitors such as aminooxyacetic acid (AOAA) (38), and BFF-122 (39), were chosen to investigate its effect on MSC-mediated cytotoxicity/growth inhibitory effects. When the inhibitors were used in combination with MSC, AOAA protected the cells from MSC-mediated toxicity. AOAA is a nonspecific transamination inhibitor, previously shown to inhibit the activity of KYAT proteins (40). A recent study presented a high-resolution crystal structure of the AOAA bound to PLP in the KYAT1 active site (41). It was evident that the small molecule inhibitor irreversibly forms an oxime with the co-factor PLP that prevents its regeneration which in turn renders KYAT1 inactivation. This corroborates our finding of protection from MSC cytotoxicity when cells were co-treated with AOAA. MSC cleavage can occur by the enzyme CTH (cystathionase) which is a γ-lyase that can exert β-elimination depending on the substrate (25). To rule out any involvement of endogenous cystathione gamma-lyase (γCTH) in the metabolism of MSC, both the γCTH inhibitor DL-propargylglycine (PAG) (42) and the substrate competitive inhibitor homoserine (HS) (43) were tested but did not show any significant efficacy in enzyme activity (Supplementary Fig. S2).

Foster *et al* showed an accumulation of Se in the pancreas, liver, and kidneys upon oral administration of ^75^Se (44). Rooseboom *et al* reported the highest level of β-elimination in the kidneys, followed by the liver, upon treatment with Se-cysteine conjugates (45). Thus, the liver represents an ideal model to validate MSC as a prodrug, with the potential advantage of adding a high-level of tumor specificity by encapsulating target mRNA in LNPs coated with ligands that mediate entry via receptors on tumor cells. We employed a microRNA strategy to reduce off-target effects. Herein, we have demonstrated that the integration of the miR122 target site to mRNA reduced the off-target expression of KYAT1 *i.e*., the addition of multiple copies (3X) of a complementary miRNA122 site can suppress KYAT1 expression in cells that have a high level of endogenous miR122. miR122 is a phenotypic determinant of normal hepatocytes (46). A similar miR122 approach was shown with apoptotic inducing proteins such as PUMA and caspase (18), but this is the first demonstration of modulation of a metabolic enzyme by this route. The use of a harmless metabolic enzyme is a great advantage compared to the induction of cell death proteins since any off-target effects would be less serious as compared to cell death proteins. This approach opens a new avenue to target and induce the expression of a protein of interest to designated transformed malignant cells and reduce off-target expression in normal cells.

MicroRNA-guided tumor-specific induction of KYAT1 in combination with the non-toxic prodrug (MSC) has a great potential in the treatment of cancers that are currently beyond cure such as HCC. A combination with factors controlling enzyme activity and tumor-specific delivery may offer further and improved therapeutic options. This concept could be extended to a variety of tumors by exploring cancer-specific ligands, antigens and miRNA combinations.

## Acknowledgments

We would like to express our gratitude to Dr. Antje Zickler from the Department of Laboratory Medicine, Karolinska Institute for cloning the KYAT1 overexpressing plasmid.

## Author contributions

M.B. and H.S conceived the idea of the study. A.K.S. performed most of the experiments and contributed to the study design and R.J. performed Fig.3A and 3C. R.G. isolated primary human normal and tumor liver hepatocytes. M.B., H.S., A.A. and A.K.S. wrote the manuscript. All authors reviewed the final version of the manuscript.

