## Supplementary Figure 1&2 and Table 1&2 for "Targeted enzyme assisted chemotherapy (TEAC) – a novel microRNA-guided and selenium-based regimen to specifically eradicate hepatocellular carcinoma"

[2]*Present address: Unit for Molecular Cell and Gene Therapy Science, Clinical Research Centre, Department of Laboratory Medicine Karolinska Institutet, Stockholm SE-141 86, Sweden.*

#### Supplementary figures 1

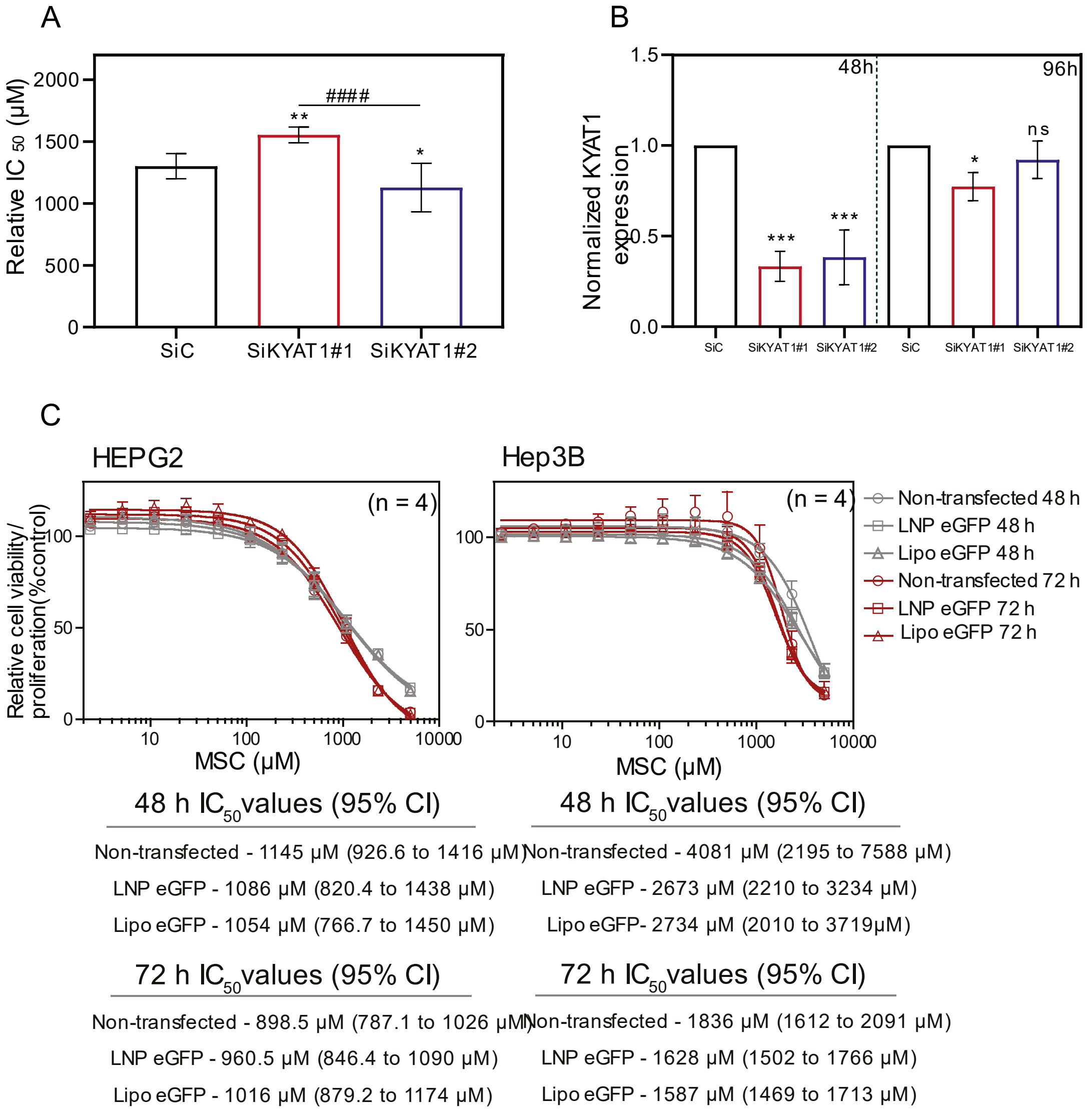


**Supplementary figure 1**. **KYAT1 knockdown protected the Huh7 cells from MSC induced cytotoxicity**. **(A)** KYAT1 siRNA mediated knockdown reverses the sensitivity of MSC mediated cytotoxicity at 72 h in Huh7 cells (n=8). The $IC_{50}$ values of KYAT1 knockdown Huh7 cells were 1302±102, 1554±65 & 1128±196 for control siRNA, KYAT1 siRNA#1 & KYAT1 siRNA#2 respectively. **(B)** Verification of knockdown efficiency of two KYAT1 siRNA (Ambion, Life Technologies, Carlsbad, CA, USA, #S2498, #S225044) at 48 h & 96 h by qRT-PCR. Negative control siRNA (Ambion, Life Technologies, Carlsbad, CA, USA, #AM4611) was used as control and KYAT1 gene expression was normalized with HPRT1 gene (n=3). Graph represent mean ± SD, statistical analysis performed with one-way ANOVA with 95% confidential interval followed by **(A)** Tukey's multiple comparison test **(B)** Dunnett's multiple comparison test (ns = not significant, * $p < 0.05$, ** $p < 0.01$, *** $p < 0.001$ and **** $p < 0.0001$ compared with SiRNA control and $^{\#}$ p<0.05, $^{\#\#}$ p<0.01, $^{\#\#\#}$ p<0.001 & $^{\#\#\#\#}$ p<0.0001 compared between two KYAT1 SiRNA). **(C) Interference of LNP's with MSC cytotoxicity.** $IC_{50}$ values of 0.5 µg/mL of eGFPmRNA (Lipo or LNP) transfected cells upon MSC treatment in HEPG2 and Hep3b cells at 48 h (n=4) and 72 h (n=4) with 95% confidential intervals. Lipo eGFP: transfected with lipofectamine 3000 (Invitrogen).

#### Supplementary figures 2

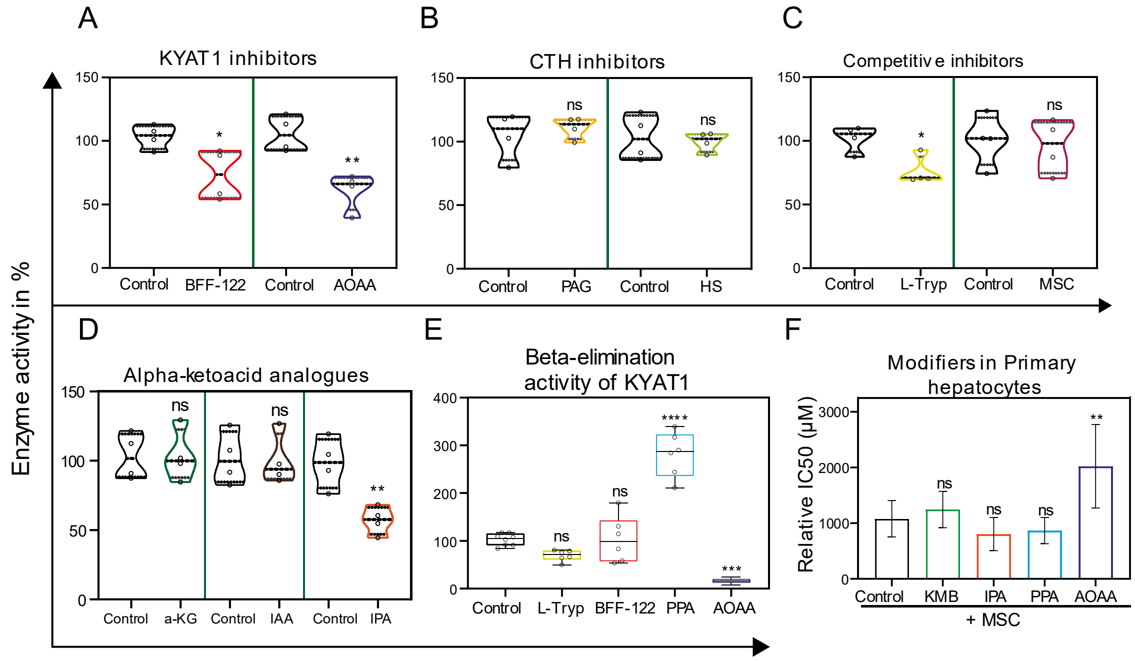


**Supplementary figure 2. KYAT 1 enzyme activity altered upon treatment with different pharmacological agents**. Transamination activity of KYAT1 was determined from cell lysates transfected with KYAT1mRNA encapsulated in LNP (0.5 µg/mL). **(A)** KYAT inhibitors (BFF-122, AOAA), (n=4). **(B)** Y-CTH inhibitors (PAG and HS), (n=4). **(C)** competitive inhibitors (L-Tryp and MSC) (n=4) and **(D)** α-Ketoacid analogs (α-KG, IAA and IPA) (n=4) relative to untreated control. **(E)** The same cell lysate was used to determine the beta-elimination activity of KYAT1 with different inhibitors and modifiers such as L-Try, BFF-122, PPA and AOAA, (n=6). **(F)** Cytotoxicity of MSC co-incubation with α-ketoacids/KYAT inhibitors (KMB, IPA, PPA and AOAA) in freshly isolated human hepatocytes at 72 h. $IC_{50}$ values for MSC, MSC+KMB, MSC+IPA, MSC+PPA and MSC+AOAA were 1079± 325, 1243± 326, 803± 298, 867± 235 and 2021± 748 respectively (n=7). **(A-F)** 20µg of protein were used in transamination and beta-elimination activity assay along with different inhibitors and inducers such as L-Try, L-Tryptophan (2.0 mM); HS, Homoserine (400 µM); PAG, Propargylglycine (1.0 mM); IAA, 3-Indoleacetic acid (1.0 mM); BFF-122 (50 µM); IPA, Indole pyruvic acid (200 µM); PPA, phenylpyruvic acid (400µM) AOAA, Aminooxyacetic acid (1.0 mM); α-KG, α-Ketoglutarate/dimethyl 2-oxoglutarate (2.0 mM); MSC, Se-methylselenocysteine (5.0 mM) and KMB, Keto-γ-(methylthio) butyric acid (100µM). Graph represent mean ± SD, statistical analysis performed with **(A-D)** Unpaired t-test **(E) & (F)** one-way ANOVA with 95% confidential interval followed by Tukey's multiple comparison test (ns = not significant, * $p < 0.05$, ** $p < 0.01$, *** $p < 0.001$ and **** $p < 0.0001$ compared with untreated Control).

#### Supplementary tables 1

**Table S1A.** Transamination activity of KYAT1 in HEPG2 cell lysate transfected with varying concentrations of eGFP and KYAT1mRNA encapsulated in LNP. Results are presented as mean ± SD from at least five independent experiments.

| Concentration in µg/mL | LNP eGFPmRNA | LNP KYAT1mRNA |
|---|---|---|
| | nmol of PPA-enol formed/min/mg of protein | |
| 0.1 | 0.35±0.29 | 9.1±1.4 |
| 0.25 | 0.34±0.3 | 15.8±4.9 |
| 0.5 | 0.53±0.42 | 18.6±3.8 |
| 1.0 | 0.45±0.19 | 23±6.8 |

**Table S1B.** Transamination activity of KYAT1 in HEPG2 cell lysate measured at different time points in cells transfected with 0.5 µg/mL of eGFP and KYAT1mRNA encapsulated in LNP. Results are presented as mean ± SD from at least four independent experiments.

| Time in hr | 0.5µg/ml of LNP eGFPmRNA | 0.5µg/ml of LNP KYAT1mRNA |
|---|---|---|
| | nmol of PPA-enol formed/min/mg of protein | |
| 24 | 0.52±0.33 | 15.1±2.9 |
| 48 | 0.7±0.51 | 18.8±5.5 |
| 72 | 0.61±0.38 | 16.8±4.0 |
| 96 | 0.73±0.22 | 16.1±4.4 |
| 120 | 0.59±0.31 | 11.1±2.6 |

### Supplementary tables 2

**Table S2.** Relative $IC_{50}$ values for MSC alone or in combination with modifiers at 72 h in HEPG2, Hep3B and Huh7 cells transfected with 0.5 µg/mL of eGFPmRNA and KYAT1mRNA encapsulated in LNP. Results are presented as mean ± SD from triplicate measurements from at least six independent experiments. Resultant $IC_{50}$ values are denoted in µM.

| Treatments | HEPG2 | | Hep3B | | Huh7 | |
|---|---|---|---|---|---|---|
| | eGFP | KYAT1 | eGFP | KYAT1 | eGFP | KYAT1 |
| MSC | 1150±170 | 472±158 | 1517±138 | 810±58 | 1223±104 | 665±219 |
| MSC+KMB | 920±282 | 346±162 | 1190±139 | 797±126 | 1010±154 | 407±213 |
| MSC+IPA | 951±212 | 414±153 | 445±48 | 286±66 | 500±129 | 107±40 |
| MSC+PPA | 1051±169 | 165±67 | 1225±192 | 269±87 | 588±123 | 107±41 |
